# SlAN1 is a limiting factor for the light-dependent anthocyanin accumulation in fruit tissues of purple tomato

**DOI:** 10.1101/2024.04.02.587792

**Authors:** Gabriel Lasmar dos Reis, Chaiane Fernandes Vaz, Luis Willian Pacheco Arge, Adolfo Luís dos Santos, Samuel Chaves-Silva, Lázaro Eustáquio Pereira Peres, Antonio Chalfun-Junior, Vagner Augusto Benedito

## Abstract

Anthocyanins are specialized plant metabolites with significant dietary value due to their anti-inflammatory properties. Research indicates that dietary intake of these phenolic compounds contributes to preventing various chronic diseases. As the most consumed vegetable worldwide, tomato (*Solanum lycopersicum*) is an excellent candidate for anthocyanin-enrichment strategies. In tomato, activation of anthocyanin biosynthesis is light-dependent, but this mechanism has yet to be entirely characterized. We investigated the role of light in anthocyanin biosynthesis in fruits of the purple tomato, which is a near-isogenic line (NIL) derived from wild accessions into cv. Micro-Tom (MT). MT-*Aft/atv/hp2* starts accumulating anthocyanin early during fruit development but is restricted to the peel (exocarp and epicarp). Manipulating light incidence in different fruit tissues determined that the absence of anthocyanin accumulation in the flesh results from the sun-blocking effect of the cyanic epicarp on the mesocarp, thus preventing light from penetrating deeper into the fruit. Transcriptional analyses of the fruit peel and flesh revealed that the bHLH transcription factor SlAN1 (Solyc09g065100) is the limiting factor for light-dependent anthocyanin accumulation in both tissues. This research enhances our comprehension of the genetic and environmental regulation of anthocyanin accumulation in fruit tissues, offering valuable insights into plant breeding for human nutrition.

**Highlight:** The expression of the *SlAN1* gene is activated in response to light signals, and it is the limiting factor for anthocyanin pigmentation in tomato fruit tissues.

## Introduction

Anthocyanins are natural pigments derived from the plant’s specialized metabolism that confer red, pink, purple, or blue pigmentation to plant tissues, depending on the molecular structure and vacuolar pH for their final hue (Hichri *et al*., 2010; Houghton *et al*., 2021). Beyond the ecological notion that anthocyanins are responsible for attracting pollinators and seed dispersers, they also play a protective role due to their antioxidant activity by scavenging reactive oxygen species (ROS) that otherwise could severely damage plant tissues (Buer *et al*., 2010; Corso *et al*., 2020). Furthermore, it has been suggested that anthocyanins form a protective barrier in plant tissues by absorbing the UV-B radiation potentially harmful to the photosynthetic machinery (Gould *et al*., 2010; Cerqueira *et al*., 2023). These properties make anthocyanins an important protective compound against environmental stresses. From a dietary perspective, based on their antioxidant and anti-inflammatory properties, anthocyanins are bioactive in preventing or mitigating a series of chronic diseases (Martin *et al*., 2011; Panchal *et al*., 2022), such as cardiovascular disorders (Cassidy *et al*., 2013), type-2 diabetes (Fallah *et al*., 2020), obesity (Muraki *et al*., 2013), and cancer (Butelli *et al*., 2008). Therefore, they represent an important health-promoting compound that should be consistently incorporated into the diet.

Considering the ubiquity of horticultural crops in the human diet, studies were carried out on these species to understand the regulatory mechanisms of the anthocyanin biosynthesis pathway (for a review, cf. Chaves-Silva *et al*., 2018). In this context, anthocyanin-enriched (cyanic) versions of fresh produce represent a critical source of anthocyanins that is readily available and cost-effective, allowing dietary enrichment simply by choosing cyanic varieties. Anthocyanins are typically found in limited quantities in cyanic products, as they are often confined to the epidermal and subepidermal cells (epicarp, exocarp, or peel), which account for only 3–5% of the fruit’s total mass (Sestari *et al*., 2014; Chaves-Silva *et al*., 2018). Thus, devising breeding strategies to generate anthocyanin-enriched varieties with an emphasis on the mesocarp (flesh) of horticultural crops is critical.

Worldwide, the tomato (*Solanum lycopersicum*) is the most consumed vegetable. It is thus an excellent model for discovering strategies aiming at anthocyanin enrichment. Although the fruits of cultivated varieties of tomato do not accumulate anthocyanins (Povero *et al*., 2011; Sestari *et al*., 2014), some related wild species, such as *S. lycopersicoides*, *S. peruvianum*, and *S. chilense*, accumulate small amounts in the subepidermal layers under adequate light conditions (Bedinger *et al*., 2011; Chaves-Silva *et al*., 2018).

Traditional breeding has delivered some varieties with cyanic tomato fruits. The introgression of the alleles *Anthocyanin fruit* (*Aft*) and *atroviolacea* (*atv*) from *S. chilense* and *S. cheesmaniae*, respectively, into *S. lycopersicum* led to purple tomato varieties, such as the cv. Indigo Rose, with an epicarp with high levels of anthocyanins (Mes *et al*., 2008; Gonzali *et al*., 2009). The further stacking of the mutation *high pigment 2* (*hp2*) from cv. Manapal, which confers hypersensitivity to light-mediated responses, into the double mutant in the cv. Micro-Tom background led to a genotype (MT-*Aft/atv/hp2*) with very high anthocyanin content in the epicarp (Sestari *et al*., 2014).

The anthocyanin biosynthesis pathway is controlled by the action of specific transcription factors (TFs), which are influenced by plant development and environmental stimuli (Albert *et al*., 2014). Among these, R2R3 MYBs can act individually or together with bHLH and WDR TFs in a multiprotein complex (MBW). On the other hand, R3 MYB are competitive inhibitors of the anthocyanin biosynthesis pathway mediated by the MBW. This complex controls the expression of the structural genes, the “*early biosynthetic genes*” (EBGs) and “*late biosynthetic genes*” (LBGs), which code for enzymes essential to the anthocyanin biosynthesis in different tissues (Chaves-Silva *et al*., 2018; Colanero *et al*., 2020*a*). In tomato, *ATV* is an R3 MYB, while *AFT* is a putative R2R3 MYB. This is consistent with the observation that the recessive *atv* allele, characterized by a loss-of-function due to a premature stop codon, and the dominant *Aft* allele both contribute to the enhancement of anthocyanin biosynthesis (Cao *et al*., 2017; Colanero *et al*., 2018; Colanero et al., 2020*b*).

Light is one of the most critical environmental factors that control the anthocyanin biosynthesis (Albert *et al*., 2009). Shading or dark conditions repress the expression of structural genes in this pathway (Hong *et al*., 2015; Liu *et al*., 2020). In some tomato genotypes (e.g., cv. ‘Indigo Rose’ and *Aft/Aft*: LA1996), the expression of anthocyanin-related genes in the epicarp starts at the mature green stage and is influenced by the amount of light received on each side of the fruit, with the shaded side exhibiting a green hue compared to a darker purple hue on the side exposed to direct light (Qiu *et al*., 2019; Colanero *et al*., 2020*b*).

A critical unresolved question is the mechanism behind internal parenchymatic tissues often less prone to accumulate anthocyanins than epidermal tissues in plants (Chaves-Silva *et al*., 2018). Here, we investigated how light influences anthocyanin accumulation in purple fruit tissues of tomato genotype MT-*Aft/atv/hp2* by blocking light during fruit development. We explored global transcriptional activity in response to light in dissected tissues of the fruit. This work sheds light on the transcriptional regulation of anthocyanin pigmentation in response to light. Our study also provides new insights for achieving higher anthocyanin content in fleshy fruits.

## Material and Methods

### Plant material, growth conditions, and sampling

To characterize the starting point of anthocyanin pigmentation in tomato fruits, we used the cyanic genotype (MT-*Aft/atv/hp2*) (Sestari *et al*., 2014) and the cultivar Micro-Tom (Meissner *et al*., 1997) as a control. The plants were grown in a greenhouse under a 16-h photoperiod with 600–700 W/m2 radiation, 21°C ± 2°C, and 50% RH. The images were analyzed using the NIS-Elements software.

Individual flowers were covered with aluminum foil at anthesis and kept for 30 days, aiming at fruit development in a complete absence of light (Supplementary Fig. S1). Subsequently, the developed fruits were exposed to light and collected at different times: immediately (0d), 2 days (2d), and 5 days (5d) after the cover was removed. For a positive control treatment (Ctl), flowers were marked at the anthesis, left exposed to normal light, and the fruits were collected after 30 days.

Young leaves and fruits in three development stages (green, turning, and mature) were collected from MT-Aft/atv/hp2 for the RT-qPCR analyses.

All fruits collected were dissected into the epicarp (the peel or exocarp) and mesocarp (the flesh), and seeds were discarded. All plant material was snap-frozen in liquid nitrogen and stored in a −80°C freezer until use. Material from three plants was collected and pooled to represent a biological replicate, and three biological replicates were used in the following analyses.

### RNA Isolation, library preparation, and sequencing

Total RNA was extracted from the epicarp and mesocarp separately using the mirVana miRNA isolation kit (Ambion, ThermoFisher) according to the manufacturer’s instructions for total RNA extraction. The integrity of the RNA was visualized on a 1.2% agarose gel, and the quantity and quality were assessed on a Nanodrop spectrophotometer. Library preparation and Illumina sequencing were performed at Novogene (Sacramento, CA, USA). Twenty-four cDNA libraries were prepared and then sequenced on an Illumina Novaseq 6000, with a configuration of 150-bp paired-end. The sequencing of transcripts revealed a total of 585.6 million reads (a mean of 23.7 million reads per library), of which 568.8 million (97%) showed sufficient quality (Q>20). The libraries were aligned on the *Solanum lycopersicum* genome assembly v.4.0 and ITAG annotation v.4.2 from SolGenomics (https://solgenomics.net; Fernandez-Pozo *et al*., 2015). On average, 22.2 million reads per library (94%) were uniquely mapped on the genome (Supplementary Table S1).

### Pre-processing and mapping of reads

Raw FastQ data were initially submitted for quality analysis with FastQC v.0.11.5 (Andrews, 2010). The cleaning step was performed with Trimmomatic v.0.39 (Bolger *et al*., 2014), with the following parameters: ILLUMINACLIP:TruSeq3-PE.fa:2:30:10:2:keepBothReads to remove adapters and keep both reads as paired, LEADING:3 and TRAILING:3 to remove bases above the quality of three at the start and final of each read, respectively, and MINLEN:36 to discard reads with < 36 bp. Each cleaned library was submitted for read mapping against the genome with the software Star v.2.7.5.c (Dobin *et al*., 2013). The counting of mapped reads for each gene was performed with featureCounts v.1.6.5 (Liao *et al*., 2014), and gene expression was normalized as TPM (transcripts per million).

### Differential expression, Functional Annotation, and Enrichment Analysis

Genes with low expression values (TPM < 1) were removed from the differential expression analysis. Differentially expressed genes were identified in R v.4.0.5 with the package DESeq2 v.1.32.0 (Love *et al*., 2014). All three replicates for each time point were used for the differential expression analysis, and the comparisons were performed against the same tissue. Genes with adjusted p-values < 0.05 were considered differentially expressed (DEGs). The package ggplot2 v.3.3.2 (Wickham, 2016) was used to build bar charts, and ComplexHeatmap v.2.9.0 (Gu *et al*., 2016) for heatmaps.

Tomato gene features were retrieved from different databases. Transcription factor information was retrieved from the PlantTFDB v.5.0 (Jin *et al*., 2016). Annotation of genes encoding metabolic enzymes was collected from the KEGG (Kyoto Encyclopedia of Genes and Genomes) database with GhostKOALA v.2.2 (Kanehisa *et al*., 2016). Gene Ontology (GO) terms were annotated with GOMAP v.1.3.4 (Wimalanathan and Lawrence-Dill, 2021) to obtain a high coverage level of genome annotation. The R’s match function was used to retrieve functional information from the genome and match it to a set of DEGs.

To investigate the primary functional annotations of the differentially expressed (DE) gene sets in each comparison, we conducted an enrichment analysis to identify overrepresented metabolic pathways, transcription factors, and GO terms in the dataset. This approach was applied for each DE profile, separated by up-and down-regulated genes. We considered features below the p-value < 0.05 threshold for metabolic pathways and transcription factors, and FDR < 0.05 for GO terms as overrepresented. Due to the high number of enriched GO terms, we computed the semantic similarity between terms with the mgoSim function from the R package GOSemSim Ver. 2.24.0 (https://doi.org/10.1007/978-1-0716-0301-7_11, https://doi.org/10.1093/bioinformatics/btq064). Following, a dimensional reduction analysis was conducted using the umap (Uniform Manifold Approximation and Projection) function from the umap Ver. 0.2.10 (https://arxiv.org/abs/1802.03426) R package. dbscan function, package dbscan Ver. 1.1-11 (https://doi.org/10.18637%2Fjss.v091.i01) was applied to identify GO terms clusterings using eps of 0.4 and minimum points of 5. Ggplot2 and ggConvexHull were used to plot umap and bubble charts, and each clustering of GO terms was labeled by the lowest FDR value.

### Structural analysis of *SlMYB-ATV* transcripts

The *SlMYB-ATV* transcript reconstruction was performed based on the data generated from the MT and MT-*Aft/atv/hp2* transcript sequences against that reported by Sun *et al*. (2020). The sequences were reconstructed with Trinity v.2.13.0 (Grabherr *et al*., 2011). Blast v.2.8.1+ (Altschul *et al*., 1990) was used to identify the *SlMYB-ATV* transcript. ORFFinder (ncbi.nlm.nih.gov/orffinder) was used to determine the ORF (Open Reading Frame) representing the transcript of interest. Subsequently, global alignment was performed for the *SlMYB-ATV* sequences from cv. Heinz, Indigo Rose (InR), Micro-Tom, and MT-*Aft/atv/hp2* with ClustalX v.2.1 (Larkin *et al*., 2007).

### RNA isolation, DNase treatment, and cDNA synthesis

Total RNA was extracted from the leaves and fruits (epicarp and mesocarp, separately) from the green, turning, and mature stages using the TRIzol^®^ reagent following the manufacturer’s instructions. Subsequently, the extracted RNA was subjected to DNase treatment (Turbo DNA-free^TM^ kit) and reverse transcribed into cDNA using the SuperScript III Reverse Transcriptase kit with oligo(dT) primers.

### Real-time PCR analysis

The RT-qPCR was performed with an ABI PRISM 7500 Real-Time (Applied Biosystems) using the SYBR Green Master Mix with the primers listed in Supplementary Table S2. β*-tubulin* (*Solyc04g081490*) and *Glyceraldehyde 3-phosphate dehydrogenase* (*GAPDH*) (*Solyc05g014470*) were used as reference genes. The relative expression was analyzed according to Pfaffl (2001).

### Statistics

Statistical analyses of the RT-qPCR data were performed using the R software (Team, 2013). The normality of variables was assessed using the Shapiro-Wilk test. Student’s t-test was applied to data with normal distribution, and the non-parametric U test (Mann-Whitney-Wilcoxon) was applied to non-normal data. All tests were used with a significance level of 95% (*P* ≤ 0.05).

## Results

### The onset of anthocyanin pigmentation in the cyanic tomato fruit (MT-*Aft/atv/hp2*)

We monitored flowering and fruit development in MT-*Aft/atv/hp2* plants and identified that anthocyanin accumulation starts right after flower senescence. As the petals fell off and the young fruit was directly exposed to light, anthocyanin accumulated in the epicarp (Fig. 1A). In contrast, fruits of the control genotype (cv. Micro-Tom) did not show visible anthocyanin pigmentation (Fig. 1B). This pattern of anthocyanin accumulation in MT-*Aft/atv/hp2* plants is independent of further fruit development but restricted mainly to the epicarp, while the mesocarp and the region under the sepals remained acyanic (Fig. 1C).

**Fig. 1.**
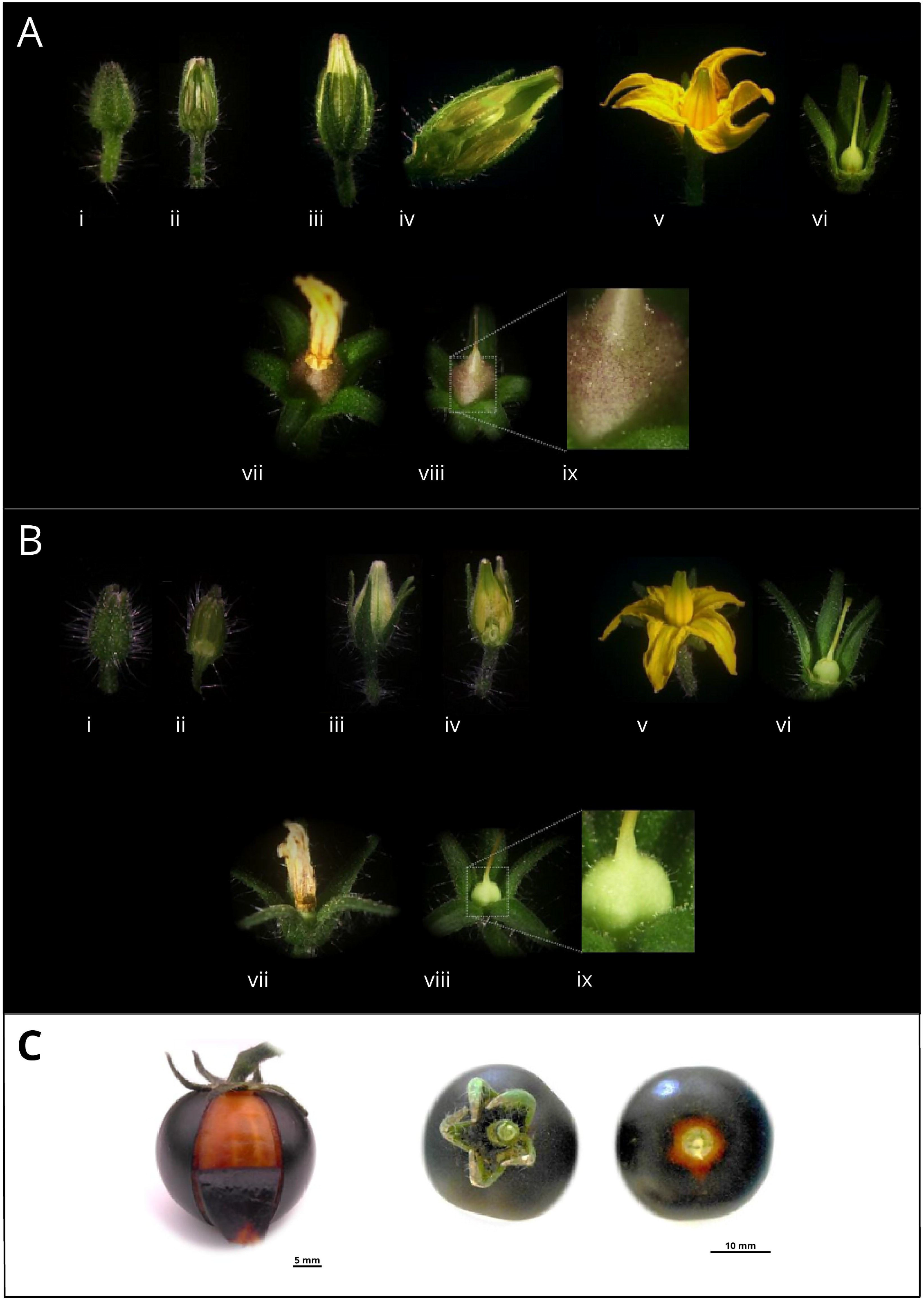
Monitoring the flowering and fruit development of two Micro-Tom (MT) genotypes and light-dependent anthocyanin accumulation patterns in purple tomato fruit. **(A)** Floral buds, flower, and developing fruit in the purple-fruit genotype, MT-*Aft/atv/hp2*, and **(B)** the regular, red-fruit cv. Micro-Tom (control). i, Developing floral bud; ii, Cross-section of the developing floral bud; iii, Immature flower; iv, Cross-section of the immature flower; v, Flower anthesis; vi, Flower anthesis without the petals; vii, Floral senescence; viii, Fruit in early development at floral senescence; ix, Zoom in on the early developing fruit shown in viii. **(C)** Lack of anthocyanin accumulation in the mesocarp and proximal region of mature fruits (MT- *Aft/atv/hp2*) when growing under normal light conditions. Notice the lack of anthocyanin accumulation in the epidermis under the calyx due to the lack of direct light exposure.

### Light activates anthocyanin biosynthesis in the tomato mesocarp

Based on the light-dependent pattern of anthocyanin pigmentation in MT-*Aft/atv/hp2* purple fruits, we investigated the metabolic response when restricting light incidence on the fruit from its first developmental stages and then exposing the physiologically mature fruit to light. Immediately after anthesis, individual flowers were covered with aluminum foil to allow the fruit to develop for 30 days to physiological maturity in the complete absence of light (Supplementary Fig. S1). As a control, flowers from other plants were marked simultaneously but not covered. Compared to control fruits, which developed a dark purple phenotype in the epicarp and a green phenotype in the mesocarp (Fig. 2), the covered fruits were completely acyanic at the removal of the cover (0d) (Fig. 2). Notably, even without light, fruits developed normally in size and shape.

**Fig. 2.**
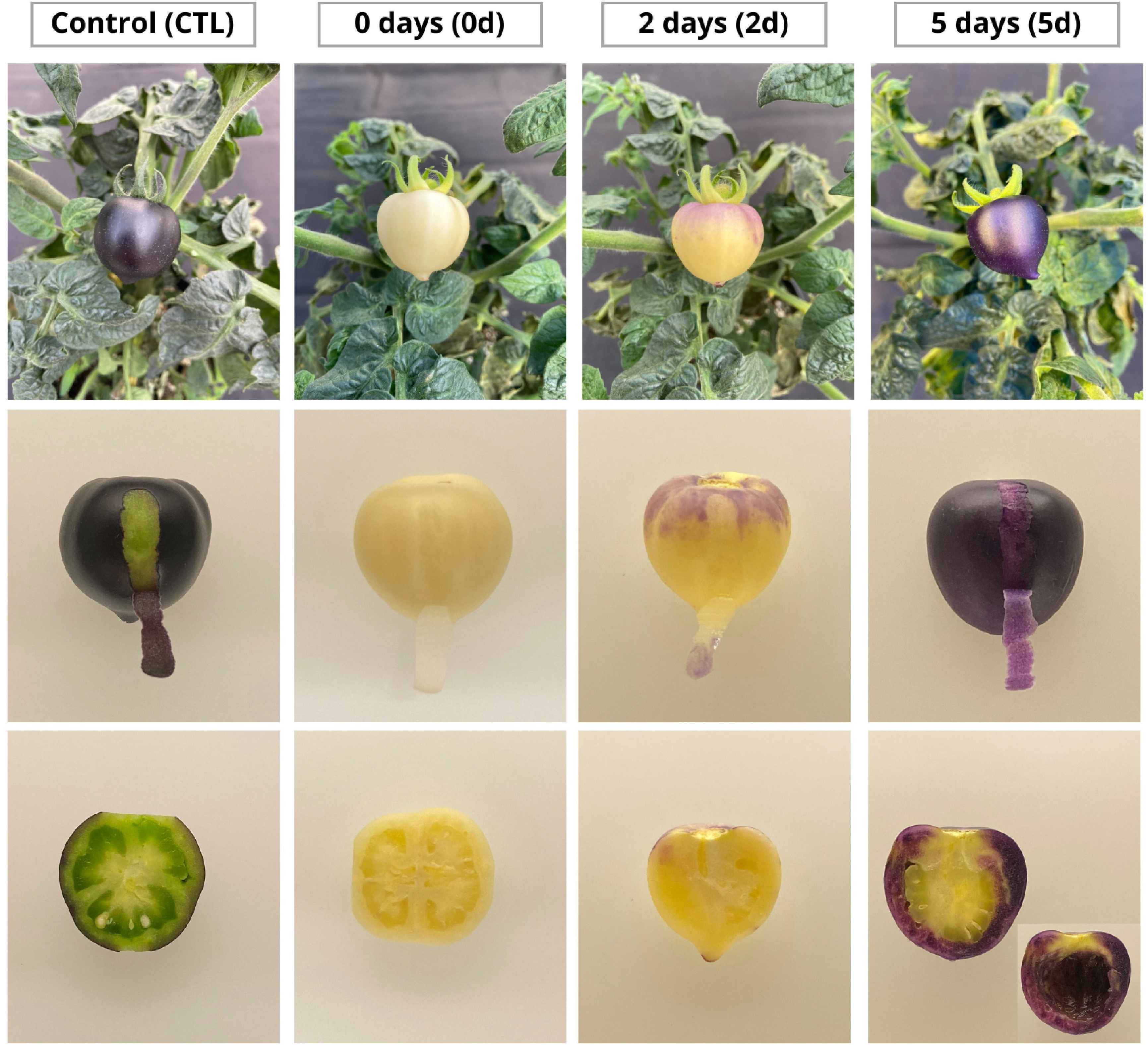
Phenotypic characterization of the anthocyanin pigmentation pattern. The epicarp and mesocarp of the cyanic tomato genotype (MT- *Aft/atv/hp2*) developed in the dark for 30 days post-anthesis. Tissues of fruit developed under different light conditions: not covered (control); immediately after the removal of the foil cover (0d); 2 days (2d); and after 5 days (5d) after cover removal. The 2d and 5d fruits were cut longitudinally to better visualize the anthocyanin accumulation in the mesocarp tissue. The inset of the 5d mesocarp cross-section displays the internal side of the mesocarp by removing the inner fruit tissues.

To investigate the activation of the anthocyanin biosynthesis after the fruit was fully developed in the dark, we exposed them to normal light conditions for up to five days after the cover had been removed and examined the anthocyanin pigmentation phenotype in the epicarp and mesocarp. Exposure to light for 2 days (2d) showed visible signs of anthocyanin accumulation in sectors of the fruit epicarp and mesocarp (Fig. 2). Furthermore, exposure to light for 5 days (5d) led to a notable increase in purple pigmentation in the entire epicarp and, surprisingly, also in the mesocarp (Fig. 2). The progressive development of the purple hue in the fruits is shown in Supplementary Fig. S2.

### Global transcriptional expression analysis of tomato fruit tissues in response to light

To further understand the molecular mechanisms behind the light-mediated transcriptional regulation of anthocyanin accumulation in the tomato fruit, we carried out RNA-seq analysis of the epicarp and mesocarp of the treatments shown in Fig. 2. By comparing the epicarp of control (fruits developed under light) and fruits exposed to light for 0 (immediately after uncovering), 2, and 5 days, we identified 4559, 5859, and 3900 differentially expressed genes (DEGs), respectively. The expression of 983 genes in the epicarp coincidently differed in 0, 2, and 5 days of light exposure compared to the control. Meanwhile, we found 1722 (0d), 5705 (2d), and 4937 (5d) DEGs in the mesocarp compared to the control (Fig. 3A; Supplementary Table S3). In this tissue, 549 genes were identified as differentially expressed coincidently in 0, 2, and 5 days of light exposure, compared to the control (Fig. 3B). The heatmap representing all DEGs is shown in Supplementary Fig. S3. Expression values for TPM (Transcript Per Million) and DEGs can be accessed in the Supplementary Table S4 and S5, respectively. This analysis revealed a light-triggered transcriptional network involved in anthocyanin accumulation in fruit tissues.

**Fig. 3.**
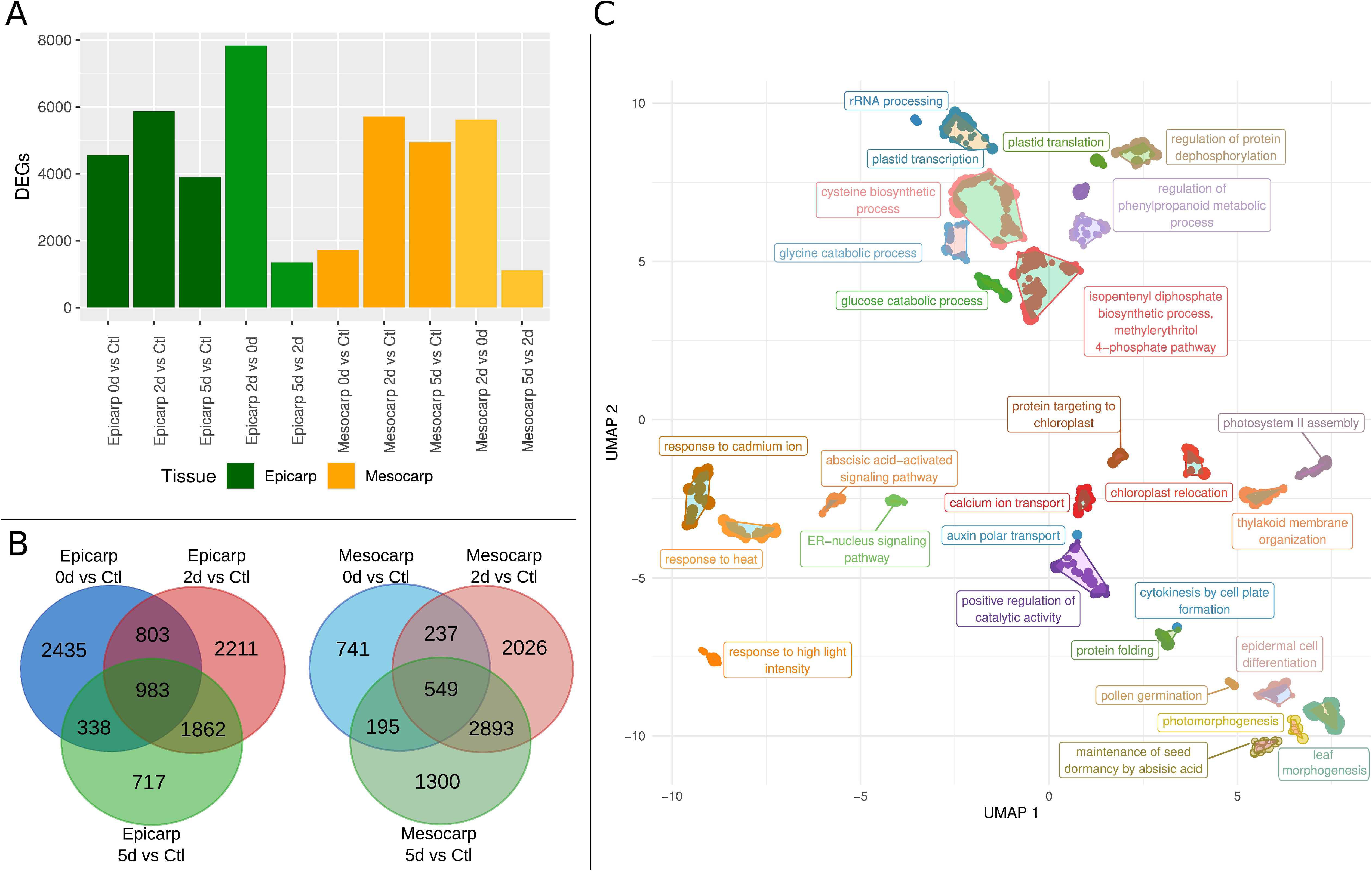
Differentially expressed genes (DEGs) and enriched GO terms in tomato fruit tissues (MT-*Aft/atv/hp2*) in response to different light exposure conditions. **(A)** Total number of DEGs in different comparisons. **(B)** Venn diagrams for differential expression within the same tissue in different light conditions versus the control (fruit grown under normal light conditions). **(C)** Summary of all enriched GO terms (biological process) clustered by semantic similarity using UMAP (Uniform Manifold Approximation and Projection). Each cluster was labeled by the GO term with the lowest FDR. d, days; Ctl, control (fruit grown under light conditions); Log2FC, logarithmic base 2 of fold change.

Enrichment analyses of functional gene annotations can reveal critical transcriptional changes associated with a phenotype. Therefore, we conducted enrichment analyses of metabolic pathways, transcription factor (TF) families, and gene ontology (GO) terms to identify distinct patterns triggered by light in fruit tissues. Seventy-five metabolic pathways were enriched in both the epicarp and mesocarp upon light exposure, including those directly or indirectly associated with anthocyanin biosynthesis and specialized metabolism: flavone and flavonol biosynthesis (map00944); flavonoid biosynthesis (map00941); phenylalanine metabolism (map00360); phenylalanine, tyrosine, and tryptophan biosynthesis (aromatic amino acids: map00400); along with the biosynthesis of carotenoids (map00906) and terpenes (map00900), and the degradation of geraniol (map00281). Aromatic amino acid biosynthesis was not enriched in the mesocarp at 0d light exposure but at 2d and 5d of light exposure when compared with the mesocarp of the control fruit developed under light conditions (Supplementary Fig. S4; Supplementary Table S6).

Transcription factors (TFs) play critical roles in modulating gene expression. We identified 15 TF families enriched in the various DEG comparisons (Supplementary Fig. S4). The most noticeable family was MYB at 2d and 5d light exposure compared to the control. Other anthocyanin-related TF families were identified, such as SQUAMOSA promoter-binding protein-like (SBP box) and Double B-box (DBB) proteins (Supplementary Fig. S4). Further analysis of the DEG sets allowed us to identify 27 clusterings of enriched GO terms associated with light-mediated responses. A clustering of light responses was found with 20 GO terms, which include response to far-red light (GO:0010218), response to red light (GO:0010114), response to high light intensity (GO:0009644), and the profiles with the highest number of enriched GO terms were down-regulated at mesocarp 2d and 5d (against control). Interestingly, anthocyanin-containing compound biosynthesis (GO:0009718) was enriched in both tissues (Supplementary Fig. S4) when anthocyanin accumulation became perceptible.

### Expression of photoreceptors genes in the anthocyanin biosynthesis pathway

We propose a model for suppressing anthocyanin accumulation in the MT-*Aft/atv/hp2* mesocarp, where the pigmented epicarp creates a shading effect on the internal tissues of the fruit, thereby preventing the mesocarp from initiating anthocyanin biosynthesis. To evaluate this hypothesis, we examined the transcriptional level of photoreceptor genes: phytochromes (sensors of red and far-red light: *SlPHYA*, *SlPHYB1*, and *SlPHYB2*); cryptochromes (sensors of blue/UV light: *SlCRY1a*, *SlCRY1b*, *SlCRY2*, and *SlCRY3*); and *SlUVR8* (a sensor of UV-B light). In our study, *SlCRY3* (*Solyc08g074270*) showed a higher expression in the epicarp than the mesocarp of fruits developed under normal light conditions (Fig. 4; Supplementary Table S7). Furthermore, *SlCRY3* expression was repressed in the epicarp of the fruit just uncovered (0d) compared to the control. In contrast, no difference was observed between the mesocarp at 0d and the control fruit (Fig. 4; Supplementary Table S7). *SlCRY3* was up-regulated in the mesocarp at 2d of light exposure compared to 0d, indicating that light reached the mesocarp and activated its expression at 2d in this tissue. On the other hand, in neither tissue, *SlCRY1a* (*Solyc04g074180*), *SlCRY1b* (*Solyc12g057040*), or *SlCRY2* (*Solyc09g090100*) showed differential expression between the epicarp and mesocarp of fruits developed under light (control), or when comparing the same tissue at 0d with the control condition (Fig. 4; Supplementary Table S7).

**Fig. 4.**
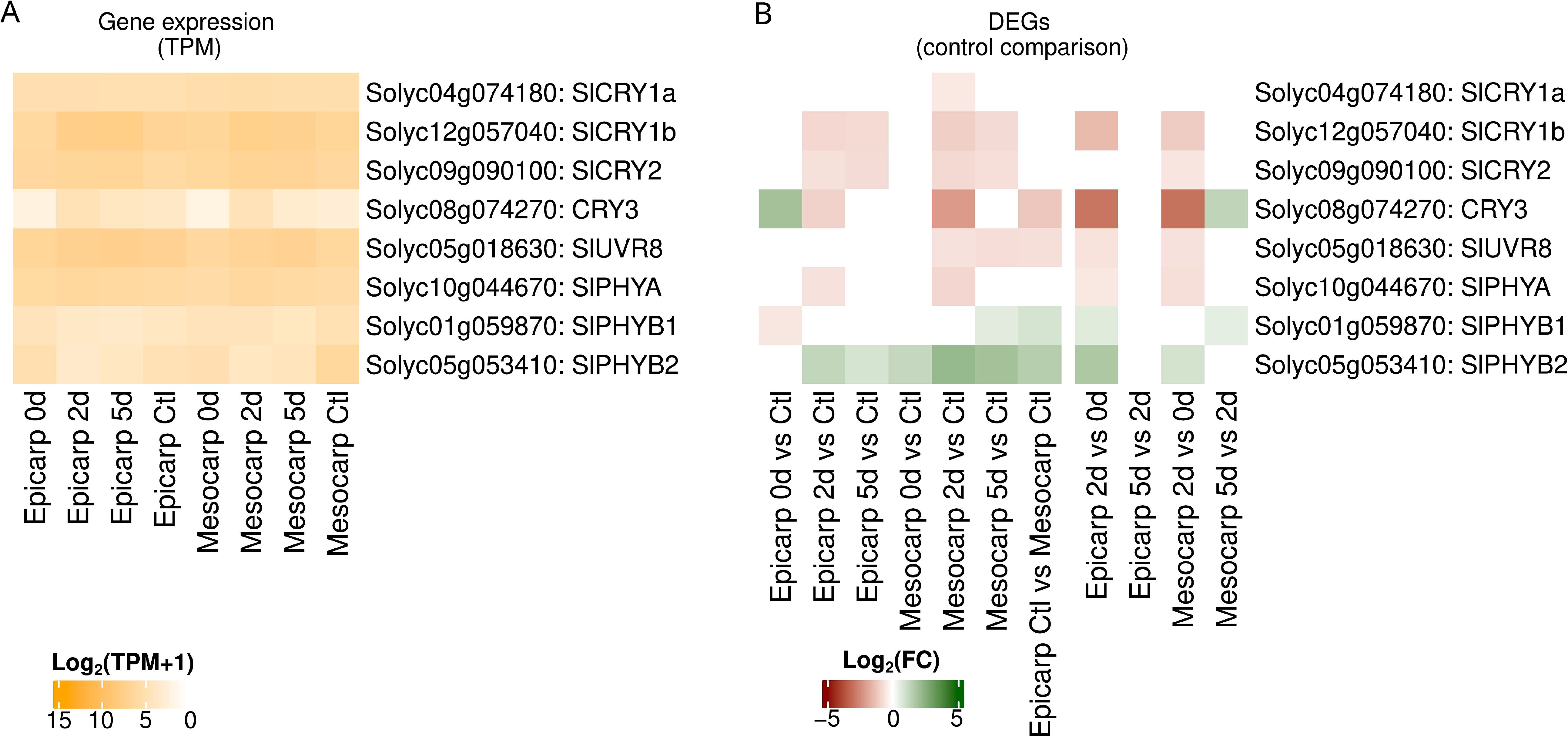
Transcriptional level of the anthocyanin photoreceptor genes in tissues of tomato fruits (MT-*Aft/atv/hp2*). Expression [log10 (TPM+1)] and differential expression (log2 FC) of the photoreceptor genes involved in the anthocyanin biosynthesis pathway. d, days of light exposure after cover removal; Ctl, control; TPM, transcripts per million; DEGs, differentially expressed genes; Log2FC, base-2 logarithm of fold change.

The UV-B-responsive photoreceptor gene *SlUVR8* (*Solyc05g018630*) showed higher expression in the epicarp compared to the mesocarp in the control condition. Furthermore, it was up-regulated in the mesocarp at 2d and 5d light exposure compared to the control mesocarp (Fig. 4; Supplementary Table S7). The expression of genes coding for the phytochromes *SlPHYB1* (*Solyc01g059870*) and *SlPHYB2* (*Solyc05g053410*) was repressed in the epicarp control compared to the mesocarp control. In contrast, *SlPHYA* (*Solyc10g044670*) did not show a significant difference in this comparison (Fig. 4; Supplementary Table S7).

### Analysis of the anthocyanin-related transcription factor genes expression in MT-*Aft/atv/hp2* tomato fruits

We analyzed the expression of some transcription factors that act upstream of the MBW complex in anthocyanin biosynthesis activation (Fig. 5; Supplementary Table S8). The expression of the *SlHY5* (*Solyc08g061130*) gene was up-regulated in the epicarp at 0d light exposure and in the mesocarp at 0d and 2d compared to the control tissues of fruits developed under normal light conditions. Interestingly, *SlHY5* expression was induced in the acyanic epicarp developed without light (0d) compared to the cyanic epicarp (control). The transcription factor *SlWRKY* (*Solyc10g084380*) was repressed in the epicarp at 0d light exposure compared to the cyanic fruit control. In contrast, it was induced in the mesocarp at 2d and 5d light exposure compared to the acyanic mesocarp of the control fruit. Furthermore, *SlWRKY* was induced in the epicarp compared to the mesocarp in control conditions. This expression pattern aligns with the development of anthocyanin pigmentation observed in the MT-*Aft/atv/hp2* fruit tissues. We also observed that the transcriptional level of *CONSTITUTIVE PHOTOMORPHOGENIC 1* (*COP1*: *Solyc05g014130*) was higher at 0d compared with 2d and 5d light exposure in both tissues, the epicarp and mesocarp (Fig. 5; Supplementary Table S8).

**Fig. 5.**
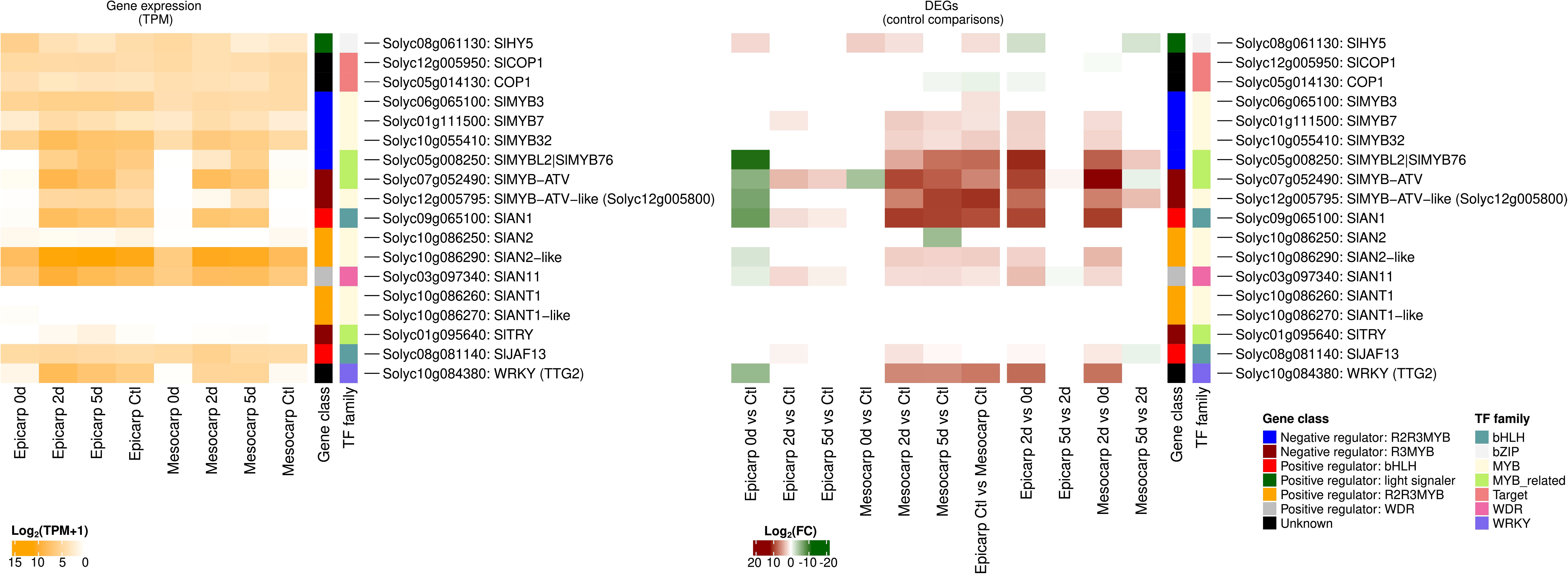
Transcriptional level of the anthocyanin biosynthetic regulatory genes in tissues of tomato fruit (MT-*Aft/atv/hp2*). Expression [log10 (TPM+1)] and differential expression (log2FC) of the anthocyanin biosynthetic regulatory genes in the epicarp and mesocarp of tomato fruits (MT-*Aft/atv/hp2*). d, days of light exposure; Ctl, control; TPM, transcripts per million; DEGs, differentially expressed genes; Log2FC, base-2 logarithm of fold change.

Among the MYB TF genes, *SlANT1* (*SlMYB113*: *Solyc10g086260*) and *SlANT1-like* (*SlMYB28*: *Solyc10g086270*) were not expressed in any of the tissues or treatments analyzed, except for the epicarp at 0d, where *SlANT1-like* showed a very low value (Supplementary Fig. S5; Supplementary Table S8). The *SlAN2* (*SlMYB75*: *Solyc10g086250*) showed very few transcripts in the epicarp and mesocarp in all conditions analyzed. By contrast, *SlAN2-like* (*SlMYB114/AFT*: *Solyc10g086290*) showed high expression levels in all tissues and conditions analyzed. In the epicarp, there was no differential expression of the *SlAN2-like* at 2d and 5d light exposure, while it was down-regulated at 0d compared to the light-exposed control. SlAN2-like expression was induced at 2d and 5d in the mesocarp but not differentially expressed at 0d compared to the control (Fig. 5; Supplementary Fig. S5; Supplementary Table S8).

SlAN2-like interacts with the constituent factors bHLH1 (SlJAF13) and WDR (SlAN11) to form the first MBW complex (Chaves-Silva *et al*., 2018). In our study, *SlJAF13* (*Solyc08g081140*) and *SlAN11* (*Solyc03g097340*) were expressed in all cyanic and acyanic tissues analyzed (Fig. 5; Supplementary Fig. S5; Supplementary Table S8). In the epicarp, *SlJAF13* expression was induced at 2d light exposure, whereas *SlAN11* expression was lower at 0d but higher at 2d and 5d when compared to the light-exposed control. In the mesocarp, *SlJAF13* and *SlAN11* expression levels were not different at 0d compared to the control but were induced at 2d and 5d. Subsequently, the formation of the first MBW complex induces the expression of bHLH2 (*SlAN1*: *Solyc09g065100*). This, in turn, replaces bHLH1 and leads to the assembly of the second MBW complex, ultimately activating the anthocyanin structural genes. In both tissues, *SlAN1* expression was minimal at 0d light exposure, whereas it highly increased at 2d and 5d (Fig. 5; Supplementary Fig. S5; Supplementary Table S8), which correlates with anthocyanin pigmentation in tomato fruit tissues.

### Expression of specific MYB regulators of anthocyanin biosynthesis in MT-*Aft/atv/hp2* tomato fruits

We examined the role of the second MBW complex in activating the expression of the negative anthocyanin regulators: *SlMYB-ATV* (*Solyc07g052490*), *SlMYBATV-like* (*Solyc12g005795*), *SlTRY* (*Solyc01g095640*), *SlMYB3* (*Solyc06g065100*), *SlMYB7* (*Solyc01g111500*), *SlMYB32* (*Solyc10g055410*), and *SlMYBL2/SlMYB76* (*Solyc05g008250*). Although these MYB repressors were expressed in all samples (Fig. 5; Supplementary Table S8), the *SlMYB-ATV* expression pattern matches that of *SlAN1*, suggesting that the second MBW complex coordinates the transcription of both genes.

Given the efficient activation of anthocyanin biosynthesis in MT-*Aft/atv/hp2* fruits, we decided to investigate if it contains a loss of function in *SlMYB-ATV*. For this, we compared the coding sequence of MT-*Aft/atv/hp2* with cv. Heinz (the reference genome), cv. Micro-Tom (the genetic background of our purple fruit genotype) and the commercial purple variety Indigo Rose. In MT-*Aft/atv/hp2*, a 4-bp insertion in the second exon of the *SlMYB-ATV* gene leads to a truncated, potentially non-functional protein without the R3 domain (Supplementary Fig. S6). The same 4-bp insertion was also observed in the Indigo Rose variety (Supplementary Fig. S6).

### Expression of structural genes associated with the anthocyanin biosynthesis in purple fruits

Regulatory and structural genes regulate the anthocyanin biosynthesis pathway (Fig. 6A). In the epicarp, the expression of almost all structural genes was highly down-regulated at 0d of light exposure, compared with the control. In contrast, at 2d and 5d of light exposure, their expression levels were similar to the control (Fig 6B; Supplementary Table S9). Interestingly, in the mesocarp at 2d and 5d light exposure conditions, EBGs and LBGs were highly up-regulated compared with the acyanic mesocarp control developed under normal light conditions, matching the anthocyanin accumulation in the fruit tissues (Fig 6B; Supplementary Table S9).

**Fig. 6.**
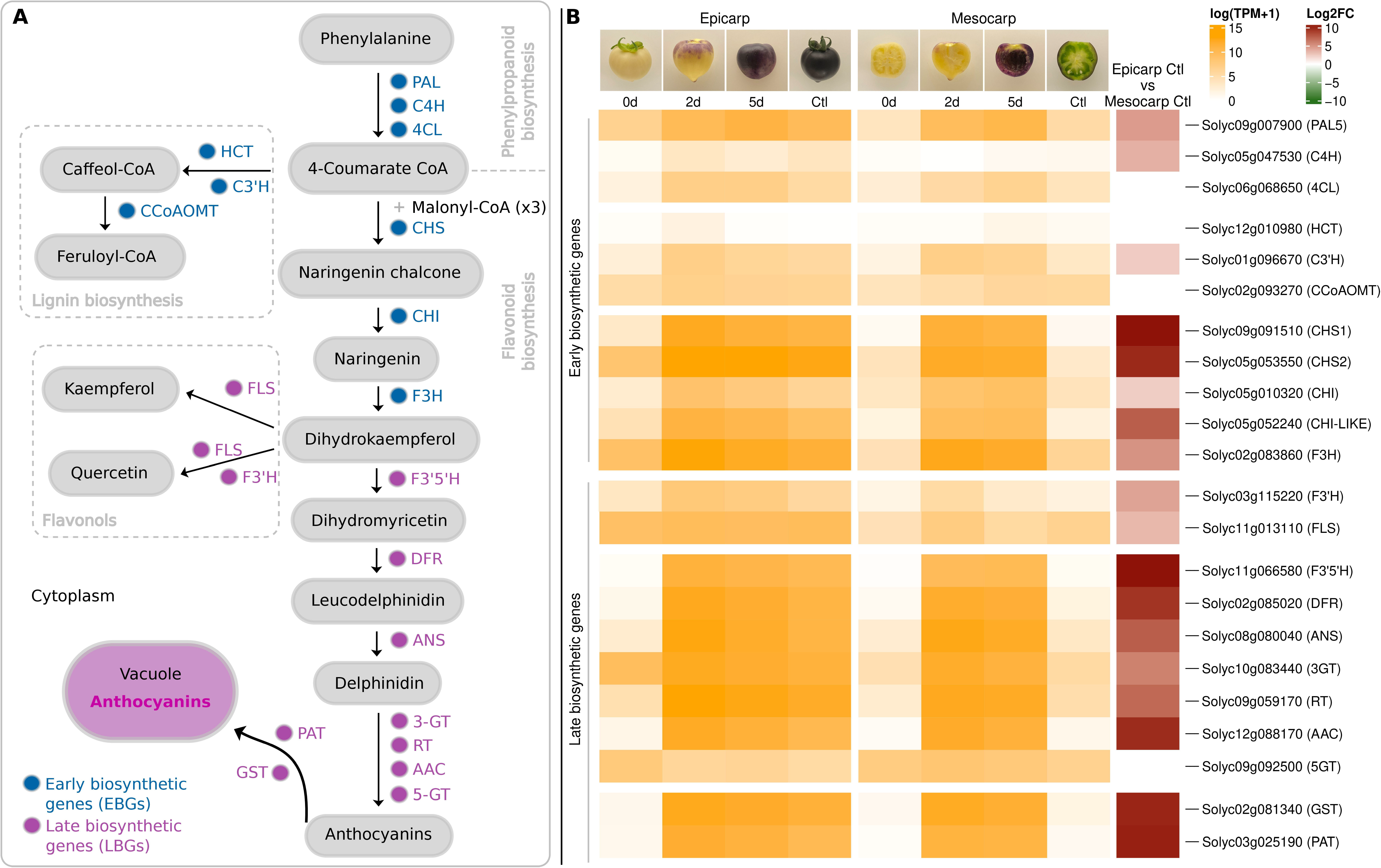
Anthocyanin biosynthesis pathway and expression patterns of the early biosynthetic genes and late biosynthetic genes in tissues of tomato fruits (MT-*Aft/atv/hp2*). **(A)** Anthocyanin biosynthesis pathway Adapted from Qiu et al. (2019). **(B)** Expression [log10 (TPM+1)] and differential gene expression (log2 FC) of the anthocyanin Early biosynthetic genes and Late biosynthetic genes in the epicarp and mesocarp of purple tomato fruits (MT- *Aft/atv/hp2*). d, days; Ctl, control; TPM, transcripts per million reads; Log2 FC, base-2 logarithm of fold change.

## Discussion

### Light-dependent anthocyanin biosynthesis activation in the MT-*Aft/atv/hp2* fruits

Genetic and environmental parameters directly influence the anthocyanin biosynthesis pathway in the tomato fruit (Albert *et al*., 2014). Light-mediated signals are usually essential to activating this pathway in different tissues (Liu *et al*., 2018*b*). In addition, fruit development also influences anthocyanin accumulation, with some anthocyanin-enriched tomato genotypes only starting to accumulate this compound after a specific stage (Qiu *et al*., 2019; Sun *et al*., 2020). Our cyanic genotype MT-*Aft/atv/hp2* is an anthocyanin-enriched tomato line that accumulates anthocyanins in the subepidermal layer of the epicarp (the peel), thus developing dark purple fruits. This line was developed by introgressing natural genetic variation from two wild species and a cultivar of tomato (the loci *Aft*, *atv*, and *hp2*) to create a near-isogenic line (NIL) in the cv. Micro-Tom background: the MT-*Aft/atv/hp2* (Sestari *et al*., 2014).

Fruits of tomato lines containing both alleles *Aft* and *atv* in homozygosity (i.e., cv. Indigo Rose) showed progressive accumulation of anthocyanins: it started right before the mature green stage on the side directly exposed to light, whereas the shaded side remained green at this stage (Qiu *et al*., 2019; Sun *et al*., 2020). On the other hand, MT-*Aft/atv/hp2* fruit started accumulating anthocyanins in the epicarp right after petal senescence (Fig. 1A). The difference in pigmentation pattern is due to the *hp2* allele, a loss of DE-ETIOLATED (DET1) function, a negative regulator of light signal transduction, conferring hypersensitivity to light (Levin *et al*., 2003). However, when growing under regular light exposure, anthocyanin accumulation in MT-Aft/atv/hp2 fruits remained restricted to the epicarp. In contrast, the mesocarp and the region under the sepals remained acyanic (Fig. 1C). This pattern shows that light is essential to activating anthocyanin biosynthesis in the purple tomato fruit.

### Anthocyanin pigmentation pattern correlates with light exposure

Anthocyanin accumulation is commonly restricted to the subepidermal cells and absent in the parenchymal cells of the mesocarp, mesophyll, and cortex. Substrate is likely to be available in these cells since the expression of specific transgenic MYB and bHLH transcription factors leads to high anthocyanin accumulation in the inner tissues of the tomato fruit (Butelli *et al*., 2008; Cerqueira *et al*., 2023). Therefore, other mechanisms should explain this “parenchymal recalcitrance”, a widespread phenomenon throughout the angiosperms (Chaves-Silva *et al*., 2018).

Here, we demonstrated that the restriction of light incidence over MT-*Aft/atv/hp2* fruits completely inhibited the anthocyanin pigmentation in all tissues (Fig. 2). Similar results of the anthocyanin pigmentation inhibition in the dark were observed in tomato (Xu et al., 2022), apple (Li *et al*., 2012), broccoli (Liu *et al*., 2020), chrysanthemum (Hong *et al*., 2015), and eggplant (Jiang *et al*., 2016; Li *et al*., 2024), confirming that light is a major factor controlling anthocyanin biosynthesis (Liu et al., 2018b). Subsequently, 5 days of light exposure of the non-pigmented, physiologically mature MT-*Aft/atv/hp2* fruits, grown in the dark for 30 days to light conditions, led to rapid activation of the anthocyanin biosynthesis in both the epicarp and mesocarp (Fig. 2). This observation shows that the activation of the anthocyanin biosynthesis pathway in the MT-*Aft/atv/hp2* fruit depends directly on the incidence of light and it is developmentally independent. Moreover, although some studies have successfully obtained anthocyanin accumulation in the mesocarp via transgenic methods (Butelli *et al*., 2008; Sun *et al*., 2020), our study is the first to report anthocyanin pigmentation in the tomato mesocarp through natural genetic variation. These findings led us to reason that the pigmented epicarp of MT-*Aft/atv/hp2* acted as a light-blocking layer starting in the first stage of fruit development, thus preventing light from reaching the mesocarp. Our experimental design, therefore, allowed the inhibition of the early pigmentation of the epicarp and facilitated the penetration of light into the non-pigmented epicarp, triggering anthocyanin biosynthesis in the mesocarp (Fig. 7).

**Fig. 7:**
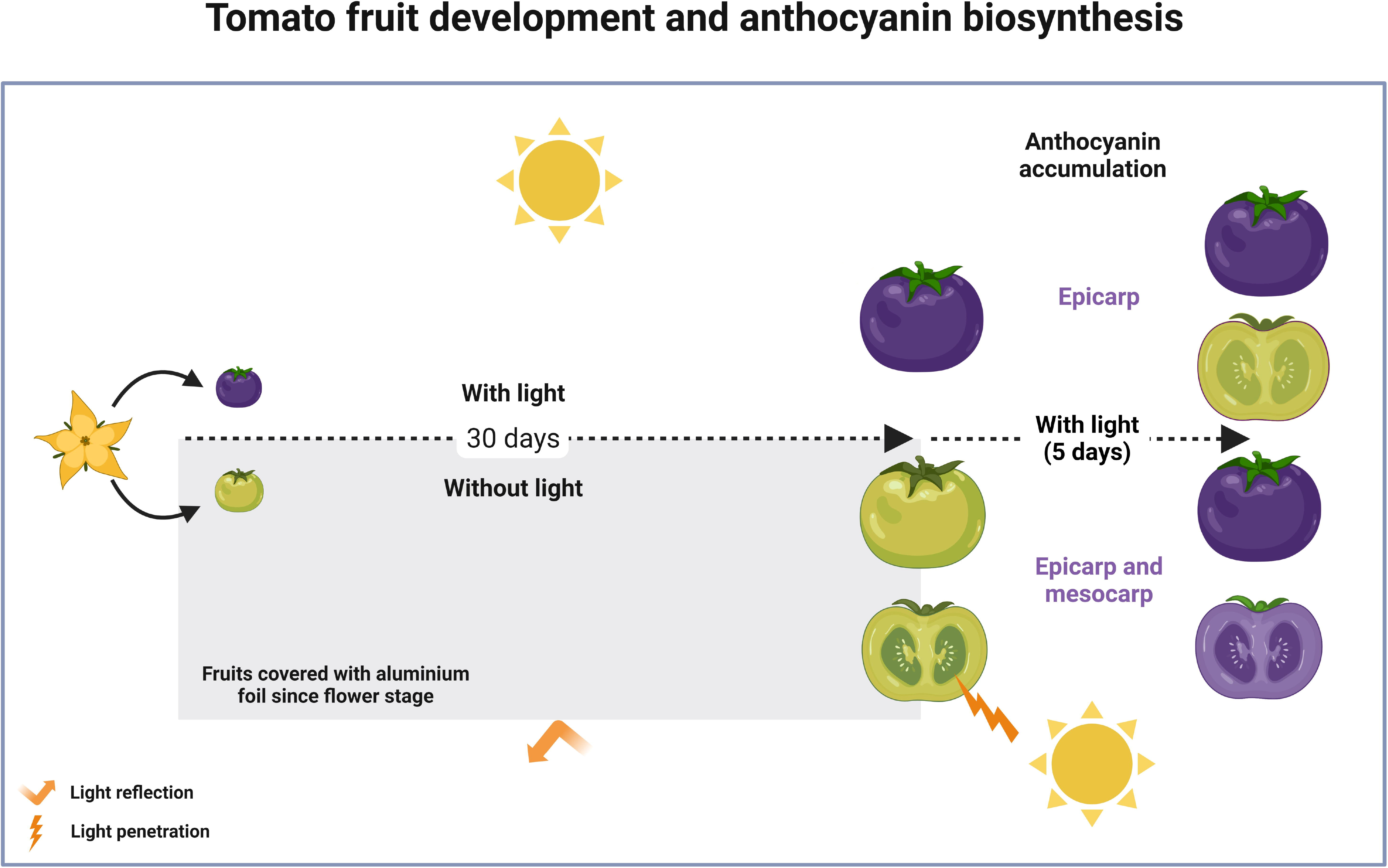
Summary of the experimental design used to manipulate light and its effect on fruit anthocyanin accumulation. At the anthesis, tomato flowers were submitted to light and dark conditions for 30 days. In light conditions, the tomato fruit showed cyanic epicarp and acyanic mesocarp. In the dark, the fruits were entirely acyanic. After the cover removal, the acyanic fruits were exposed to light for 5 days, leading to anthocyanin accumulation in the epicarp and mesocarp of these fruits.

### Photoreceptors involved in the anthocyanin biosynthesis activation in tomato fruits

Plants use specific photoreceptor classes to receive light signals and coordinate stimulus responses. The cryptochrome *CRY3* participates in anthocyanin biosynthesis in eggplant, purple broccoli, and petunia. *CRY3* expression is repressed in shading conditions and highly induced by light Fields(Li *et al*., 2017; Fu *et al*., 2020; Liu *et al*., 2020). In petunia, while *CRY3* was repressed by exposure to red light compared to white and blue lights, *CRY1* and *CRY2* did not show significant changes in response to the light quality (Fu *et al*., 2020). The synergistic effect of blue and UV-B light promoted anthocyanin accumulation in the epicarp of the tomato *Aft* line (Kim *et al*., 2021). In our study, *SlCRY3* and *SlUVR8* were the only photoreceptors transcriptionally induced in the cyanic epicarp compared with the acyanic mesocarp in control conditions. *SlUVR8* expression increased in both tissues along the days of light exposure for the fruits grown in the dark, whereas *SlCRY3* peaked at 2d when anthocyanin accumulation started becoming noticeable (Fig. 4). Based on these gene expression patterns and the current understanding that short-wavelength radiation (*i.e.* blue and UV lights) promotes anthocyanin accumulation, we infer that *SlCRY3* and *SlUVR8* are the primary photoreceptors activating the anthocyanin pigmentation in tomato fruits.

Therefore, the anthocyanin pigmentation of the MT-*Aft/atv/hp2* epicarp starts at the first stage of fruit development and directly influences the quantity and quality of light at the mesocarp by blocking short-wavelength radiation. This light-blocking effect leads to the inactivation of light-induced signal transduction in the mesocarp, which is necessary to activate anthocyanin biosynthesis-responsive genes (Fig. 7).

### COP1 may act as a negative regulator of SlHY5 activity in the tomato fruit growing in the dark

Anthocyanin biosynthesis is controlled by complex molecular mechanisms orchestrated by transcription factors and regulated by developmental and environmental stimuli. Light-responsive transcription factors can act individually or in multiprotein complexes to regulate the expression of anthocyanin regulatory and structural genes (Qiu *et al*., 2016, 2019). Upon photoreceptor perception of blue and UV-B light, anthocyanin biosynthesis relies on a signal transduction cascade coordinated by the transcription factors HY5 and COP1 (Podolec and Ulm, 2018). HY5 is a bZIP transcription factor considered a central regulator of anthocyanin enrichment in tomato fruits (Liu *et al*., 2018*a*). In the cultivar ‘Indigo Rose’, SlHY5 is related to the anthocyanin biosynthesis (Qiu *et al*., 2019; Sun *et al*., 2020); however, in the cv. Indigo Rose, the expression of SlHY5 was not significantly different between the fruit side facing the light, which developed a purple phenotype, compared to the non-cyanic epicarp on the shaded side (Qiu *et al*., 2019). Similarly, in our study, SlHY5 expression was up-regulated in the epicarp developed in the dark (0d) compared to the control cyanic epicarp (Fig. 5).

This peculiar SlHY5 expression pattern, i.e., expression in both light and dark conditions but not leading to the anthocyanin accumulation in dark conditions, could be explained by a post-translational regulation of the SlHY5 protein affecting its stability and activity, as observed in Arabidopsis (Hardtke *et al*., 2000). In the dark, COP1 ubiquitinates HY5, leading to its degradation by the proteasome. Conversely, in light conditions, photoreceptors become the targets of COP1 instead of HY5 (Saijo *et al*., 2003). Our data indicate that SlHY5, COP1, SlCRY3, and SlUVR8 expression pattern is correlated with the anthocyanin accumulation in tomato fruits. In summary, even with higher gene expression in the dark (acyanic tissues), SlHY5 may be post-translationally repressed by COP1. On the other hand, when the fruit tissues received light, SlCRY3 and SlUVR8 transcription levels were induced (Fig. 4). Furthermore, our GO term analysis showed enrichment in protein ubiquitination (GO:0016567) at 0d in the acyanic epicarp compared with the fruit growing in normal light conditions, as well as in the epicarp versus mesocarp of the control (Supplementary Fig. S4). Nevertheless, in a comparative analysis of the differential expression of the genes *SlHY5* and *COP1* between the epicarp and mesocarp, both in the control condition, *SlHY5* was up-regulated, whereas *COP1* was down-regulated in the cyanic epicarp (Fig. 5). This opposite expression pattern corroborates the hypothesis that COP1 negatively regulates SlHY5 in tomato fruits.

Transcription factors from the WRKY family are essential regulators of anthocyanin biosynthesis in an HY5-independent manner (Qiu *et al*., 2019). WRKY physically interacts with WD to form a transcriptional complex independent of the MBW complex, leading to the transcriptional activation of membrane transporter and vacuolar acidification genes (Lloyd *et al*., 2017). In our study, the *SlWRKY* transcriptional profile matches the anthocyanin accumulation pattern in MT-*Aft/atv/hp2* fruit tissues (Fig. 5). Thus, our model suggests that *SlWRKY* expression is regulated by light and that it plays an essential cyanogenic role in the tomato fruit by activating the transcription of anthocyanin structural genes.

### Transcriptional patterns of anthocyanin-positive regulatory genes may explain the lack of anthocyanin synthesis in the mesocarp

We demonstrated that anthocyanin accumulation in MT-*Aft*/*atv*/*hp2* fruits is light-dependent. Tomato lines with the dominant locus *Aft* (*Anthocyanin fruit*) display a purple phenotype linked to a genomic region that contains four in-tandem R2R3 MYB genes (Sapir *et al*., 2008; Cao *et al*., 2017): *SlAN2* (*SlMYB75*: *Solyc10g086250*), *SlANT1* (*SlMYB113*: *Solyc10g086260*), *SlANT1-like* (*SlMYB28*: *Solyc10g086270*), and *SlAN2-like* (*SlMYB114*: *Solyc10g086290*), which corresponds to the *AFT* gene. Fig. 5 and Supplementary Fig. S5 show that *SlANT1*, *SlANT1-like*, and *SlAN2* displayed no or very low expression levels in both cyanic and acyanic fruit tissues. Similar studies on tomato genotypes with the *Aft* locus found insignificant expression levels for these genes (Qiu *et al*., 2019; Colanero *et al*., 2020*b*; Sun *et al*., 2020), suggesting they are not involved in regulating anthocyanin biosynthesis in *Aft*-bearing purple tomato fruits.

In contrast, *SlAN2-like* was highly expressed in all tissues and conditions analyzed. Its expression was only slightly lower in the non-pigmented epicarp at 0d of exposure to light compared to the purple epicarp of fruits developed under normal light conditions (Fig. 5; Supplementary Fig. S5). The same pattern was reported for the ‘Indigo Rose’ cultivar and a mutant with a reduced anthocyanin pigmentation (Qiu *et al*., 2019). Also, the *SlHY5* transcription occurred even in dark conditions (Fig. 5). These findings led us to speculate about a possible negative regulation of the SlAN2-like protein by COP1 in tomato fruit tissues under dark conditions, similar to what is observed in apple and eggplant (Li *et al*., 2012, 2024).

The MYB transcription factor SlAN2-like interacts with the constitutive factors bHLH1 (SlJAF13) and WDR (SlAN11) to form the first MBW complex, which induces bHLH2 (*SlAN1*) expression. Subsequently, bHLH2 (SlAN1) replaces bHLH1 (SlJAF13) to configure the second MBW complex. This second complex, in turn, activates the expression of *SlAN1* (“reinforcement mechanism”) and the late anthocyanin biosynthetic genes (Colanero *et al*., 2020*b*). Even though *SlJAF13* and *SlAN11* expression levels fluctuated across samples, they were constitutively detected in pigmented and non-pigmented tissues (Fig. 5), confirming previous reports (Gao *et al*., 2018). Transcriptional analysis in transgenic lines and cv. ‘Indigo Rose’ purple fruit tissues showed that *SlAN1* expression correlated with the level of anthocyanin pigmentation (Butelli *et al*., 2008; Bassolino *et al*., 2013; Qiu *et al*., 2016, 2019), which was confirmed in our study (Fig. 5; Supplementary Fig. S5). Although SlAN2-like was the most highly expressed anthocyanin-related MYB factor across all tissues and conditions analyzed, its expression levels remained relatively stable in both cyanic and non-cyanic tissues of tomato fruits at 30-35 days after anthesis. Even though SlAN2-like is actively expressed in dark conditions, the protein may be inactive due to an undetermined post-translational mechanism.

The interaction between COP1 and MYB proteins controlling the anthocyanin accumulation was observed in apple and eggplant. MdCOP1 interacts with MdMYB1 to regulate light-induced anthocyanin biosynthesis in apple (Li *et al*., 2012), and SmCOP1 interacts with SmMBY5 to trigger the degradation of the latter via the 26S proteasome pathway in eggplant (Li *et al*., 2024). The expression patterns of COP1 and SlAN2-like genes observed here indicate that the interaction between their products also occurs in tomato fruits and directly influences anthocyanin accumulation. In dark conditions, COP1 may act as a negative regulator of the SlAN2-like protein, thus inhibiting the formation of the MBW complex and, consequently, SlAN1 expression. In turn, light induces the expression of photoreceptor genes, making them preferred targets of COP1 for ubiquitination instead of SlAN2-like. This mechanism allows the formation of the MBW (SlAN2-like/SlJAF13/SlAN11) complex to activate the SlAN1 expression (Fig. 8).

**Fig. 8:**
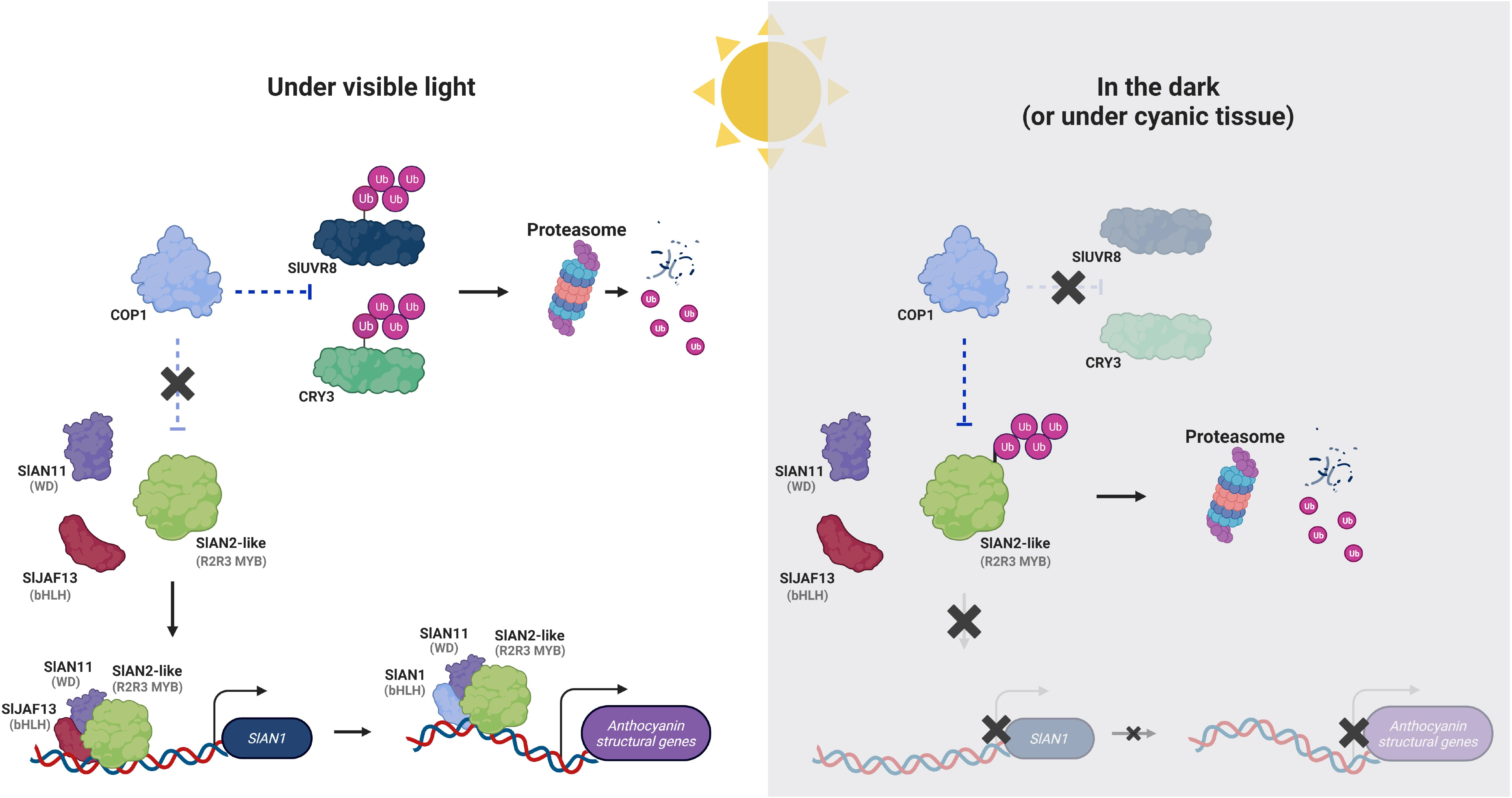
Possible transcriptional model for anthocyanin biosynthesis regulation under light and dark conditions. Under visible light, instead of ubiquitinating the SlAN2-like (*Solyc10g086290*), the COP1 (*Solyc05g014130*) ubiquitinates the photoreceptors SlUVR8 (*Solyc05g018630*) and CRY3 (*Solyc08g074270*), leading to their degradation by the proteasome. In this condition, the SlAN2- like forms the first MBW complex together with the SlJAF13 (*Solyc08g081140*) and SlAN11 (*Solyc03g097340*) to activate the *SlAN1* (*Solyc09g065100*) gene expression. After that, the SlAN1 replaces the SlJAF13 to form the second MBW complex to activate the anthocyanin structural genes. In dark conditions, the COP1 will ubiquitinate the SlAN2-like, inducing its degradation by the proteasome, inhibiting the formation of the first MBW complex from activating the *SlAN1* expression and, consequently, the anthocyanin biosynthesis.

Our findings show that *SlAN1* is the limiting factor governing the development of anthocyanin pigmentation of tomato fruit tissues, with its expression being light-dependent. As a result, the early onset of epicarp pigmentation, which obstructs light from penetrating the inner tissues, ultimately impedes *SlAN1* expression and anthocyanin accumulation in the mesocarp of MT-*Aft/atv/hp2*. This regulation may be the key to unleashing light-independent anthocyanin biosynthesis in different edible plant parts, such as purple-fleshed roots, tubers, and internal fruit organs.

### Loss-of-function of *SlMYB-ATV* and its expression in the MT-*Aft/atv/hp2* genotype

Genes that encode enzymes of the anthocyanin pathway are divided into “*early biosynthetic genes*” (EBGs) and “*late biosynthetic genes*” (LBGs) (Quattrocchio *et al*., 2006). EBGs are related to synthesizing flavonoid precursors and final products (e.g., chalcones, dihydroflavonols, and flavonols). In contrast, LBGs are more specific for anthocyanins (Fig. 6A). In our study, the expression pattern of anthocyanin structural and regulatory genes correlated with anthocyanin pigmentation in the tissues of tomato fruits (Fig. 6B).

In addition to activating *SlAN1* and LBGs, the second MBW complex also induces the expression of MYB repressors (Albert *et al*., 2014; Colanero *et al*., 2020*b*). The R3-MYB protein can directly bind to the bHLH factors, inhibiting the formation of the MBW complex activator of anthocyanin-related genes (Colanero *et al*., 2018; Sun *et al*., 2020).

*SlMYB-ATV* (*Solyc07g052490*) is responsible for negatively regulating the anthocyanin biosynthesis in tomato fruits (Cao *et al*., 2017; Colanero *et al*., 2018). The transcriptional activation of *SlMYB-ATV* is triggered by the MBW ternary complex (SlAN2-like/SlAN1/SlAN11), which also activates *SlAN1* expression (Colanero *et al*., 2018). In our study, the expression pattern of *SlMYB-ATV* matches that of *SlAN1* (Fig. 5), suggesting the activation of both genes by the same complex. The functional, dominant *SlMYB-ATV* allele is present in *S. lycopersicum*, whereas the recessive *atv* allele is present in some wild species, such as *S. cheesmaniae*. The *atv* alleles code for a truncated, non-functional protein due to a 4-bp insertion in the second exon that leads to a frameshift and the onset of a premature stop codon. The resulting truncated SlMYB-ATV protein lacks the R3-domain that cannot bind to the bHLH factors and thus cannot disrupt the MBW complex (Cao *et al*., 2017; Colanero *et al*., 2018; Sun *et al*., 2020), favoring the anthocyanin enrichment in tomato tissues. We identified that the *SlMYB-ATV* gene from the cyanic line MT-*Aft/atv/hp2* also contains the 4-bp insertion. Therefore, the nonfunctional *SlMYB-ATV* allele favors the anthocyanin pigmentation in the tissues of the tomato fruits.

### Additional traits observed in MT-*Aft/atv/hp2* plants

The yield of MT-*Aft/atv/hp2* plants was reported to be comparable to that of cv. MT. MT- *Aft/atv/hp2* fruits showed significantly higher levels of ascorbic acid, lycopene, and β-carotene than cv. MT (Sestari *et al*., 2014). Anthocyanins have also been associated with increased resistance to biotic stresses in mango (Sivankalyani *et al*., 2016). We observed that MT- *Aft/atv/hp2* plants appeared more resistant to thrips (*Thysanoptera*) than MT plants in greenhouse conditions. Even though we have not addressed this question in our current research, it warrants further investigation.

### Conclusions

This study brings novel information on anthocyanin metabolism in the mesocarp cells of purple tomato fruits mediated by light. Short wavelengths, such as blue and UV-B light, represent crucial signals for anthocyanin biosynthesis activation in cyanic tomato fruits. In the epicarp of MT-*Aft/atv/hp2*, these wavelengths are detected early during development and trigger signal transduction pathways, thereby inducing the expression of the anthocyanin regulatory and structural genes, resulting in anthocyanin accumulation. This pigmentation establishes a filter that blocks the short wavelengths from reaching the deep tissues of the fruit, consequently suppressing the expression of the *SlAN1*, the limiting gene for anthocyanin accumulation in mesocarp cells. Exposing acyanic fruits, which develop in the dark, to light for 5 days promotes anthocyanin accumulation in the epicarp and the mesocarp tissues. Therefore, our results present a working hypothesis to elucidate the anthocyanin recalcitrance observed in parenchymatic cells of inner fruit tissues. To overcome this resistance, we propose a reliable approach or genetic pathway- namely, preventing early epidermis pigmentation.

## Supplementary Data

**Supplementary Table S1.** Summary of cleaning, mapping, and counting reads on the tomato reference genome.

**Supplementary Table S2.** Primers used in the RT-qPCR analyses.

**Supplementary Table S3.** Differentially expressed genes by log2FC interval.

**Supplementary Table S4.** Gene expression in TPM (Transcript Per Million) of epicarp and mesocarp tissues from tomato fruits (MT-*Aft/atv/hp2*) developed under dark and light conditions.

**Supplementary Table S5.** Differentially expressed genes identified for different comparisons of epicarp and mesocarp tissues, FDR < 0.05.

**Supplementary Table S6.** Biological processes enriched in each profile in up- and down- regulated genes.

**Supplementary Table S7.** Expression of the photoreceptor genes in the epicarp and mesocarp of tomato fruits (MT-*Aft/atv/hp2*).

**Supplementary Table S8.** Expression [log (TPM+1)] and differential expression (log2FC) of the anthocyanin biosynthetic regulatory genes in the epicarp and mesocarp of tomato fruits (MT- *Aft/atv/hp2*).

**Supplementary Table S9.** Expression [log (TPM+1)] and differential expression (log2FC) of the anthocyanin biosynthetic structural genes in the epicarp and mesocarp of tomato fruits (MT- *Aft/atv/hp2*).

**Supplementary Fig. S1.** Restriction of light incidence during fruit development. Individual flowers were covered with aluminum foil at anthesis **(A)** for 30 days **(B).**

**Supplementary Fig. S2.** Characterization of the progressive development of anthocyanin pigmentation in the epicarp of MT-*Aft/atv/hp2* under different light conditions for 30 days after anthesis. **(A)** Thirty-day fruit developed in the dark immediately after cover removal. **(B-F)** Covered fruit exposed to normal light conditions for 1**(B)**, 2**(C)**, 3**(D)**, 4**(E)**, and 5**(F)** days after cover removal. (G) Fruit developed in normal light conditions (not covered).

**Supplementary Fig. S3.** Heatmap of DEGs. Each column compares two conditions, whereas each line represents a gene. See Table S3 for details. d, days; Ctl, control (fruit grown under light conditions); Log2FC, logarithmic base 2 of fold change.

**Supplementary Fig. S4.** Analyses of Gene Ontology (GO) terms, KEGG pathways, and transcription factors enriched in the differentially expressed gene (DEG) set within each tissue at a specific light exposure time compared to the control condition (fruit developed under regular light exposure). The last comparison shows DEG regulation between the epicarp and mesocarp in the control fruit. Ctl, control; d, days of exposure to light after cover removal.

**Supplementary Fig. S5.** qPCR analysis of regulatory *R2R3 MYB* (*SlAN2-like*, *SlAN2*, *SlANT1- like*, and *SlANT1*), *bHLH* (*SlAN11*), *WDR* (*SlAN11*), *Early biosynthetic gene* (*CHI-like*), and *Late biosynthetic gene* (*DFR*) genes performed in the young leaves and fruits (peel and flesh) of MT-*Aft/atv/hp2* at green, turning, and mature stages. Data are means of three biological replicates. Student’s T-test was performed. Different letters indicate significant differences at P ≤ 0.05.

**Supplementary Fig. S6. (A)** Sequence alignment of *SlMYB-ATV* (*Solyc07g052490*) transcripts from cv. ‘Indigo Rose’ (InR), MT-*Aft/atv/hp2*, Micro-Tom, and Heinz (genome) tomato genotypes. **(B)** Schematic representation of the *SlMYB-ATV* (*Solyc07g052490*) gene mutation in the MT-*Aft/atv/hp2*.

**Supplementary Fig. S7.** Principal component analysis based on expression values.

## Author Contributions

GLR, ALS, LEPP, AC-Jr., and VAB conceived and planned the study; GLR, ALS, and VAB designed the experiments; GLR, SC-S, and ALS performed the experiments; GLR, CFV, LWPA, and VAB analyzed the data; GLR, CFV, and LWPA wrote the manuscript; VAB, LEPP, and AC-Jr revised the manuscript; VAB supervised all steps of the study and final manuscript. All authors read and approved the manuscript.

## Data availability

The RNA-seq data underlying this article are available in the Gene Expression Omnibus (GEO) Database (GSE235565).

## References

Albert NW, Davies KM, Lewis DH, Zhang H, Montefiori M, Brendolise C, Boase MR, Ngo H, Jameson PE, Schwinn KE. 2014. A Conserved Network of Transcriptional Activators and Repressors Regulates Anthocyanin Pigmentation in Eudicots. The Plant Cell 26, 962–980.

Albert NW, Lewis DH, Zhang H, Irving LJ, Jameson PE, Davies KM. 2009. Light-induced vegetative anthocyanin pigmentation in Petunia. Journal of Experimental Botany 60, 2191–2202.

Altschul SF, Gish W, Miller W, Myers EW, Lipman DJ. 1990. Basic local alignment search tool. Journal of Molecular Biology 215, 403–410.

Andrews S. 2010.FastQC: a quality control tool for high throughput sequence data.

Bassolino L, Zhang Y, Schoonbeek HJ, Kiferle C, Perata P, Martin C. 2013. Accumulation of anthocyanins in tomato skin extends shelf life. New Phytologist 200, 650–655.

Bedinger PA, Chetelat RT, McClure B, et al. 2011. Interspecific reproductive barriers in the tomato clade: Opportunities to decipher mechanisms of reproductive isolation. Sexual Plant Reproduction 24, 171–187.

Bolger AM, Lohse M, Usadel B. 2014. Trimmomatic: A flexible trimmer for Illumina sequence data. Bioinformatics 30, 2114–2120.

Buer CS, Imin N, Djordjevic MA. 2010. Flavonoids: New roles for old molecules. Journal of Integrative Plant Biology 52, 98–111.

Butelli E, Titta L, Giorgio M, et al. 2008. Enrichment of tomato fruit with health-promoting anthocyanins by expression of select transcription factors. Nature Biotechnology 26, 1301–1308.

Cao X, Qiu Z, Wang X, et al. 2017. A putative R3 MYB repressor is the candidate gene underlying *atroviolacium*, a locus for anthocyanin pigmentation in tomato fruit. Journal of Experimental Botany 68, 5745–5758.

Cassidy A, Mukamal KJ, Liu L, Franz M, Eliassen AH, Rimm EB. 2013. High anthocyanin intake is associated with a reduced risk of myocardial infarction in young and middle-aged women. Circulation 127, 188–196.

Cerqueira JVA, Zhu F, Mendes K, Nunes-Nesi A, Martins SCV, Benedito VA, Fernie AR, Zsögön A. 2023. Promoter replacement of *ANT1* induces anthocyanin accumulation and triggers the shade avoidance response through developmental, physiological and metabolic reprogramming in tomato. Horticulture Research 10, uhac254.

Chaves-Silva S, Santos AL dos, Chalfun-Júnior A, Zhao J, Peres LEP, Benedito VA. 2018. Understanding the genetic regulation of anthocyanin biosynthesis in plants – Tools for breeding purple varieties of fruits and vegetables. Phytochemistry 153, 11–27.

Colanero S, Perata P, Gonzali S. 2018. The *atroviolacea* gene encodes an R3-MYB protein repressing anthocyanin synthesis in tomato plants. Frontiers in Plant Science 9, 1–17.

Colanero S, Perata P, Gonzali S. 2020a. What’s behind purple tomatoes? Insight into the mechanisms of anthocyanin synthesis in tomato fruits. Plant Physiology 182, 1841–1853.

Colanero S, Tagliani A, Perata P, Gonzali S. 2020b. Alternative Splicing in the *Anthocyanin Fruit* Gene Encoding an R2R3 MYB Transcription Factor Affects Anthocyanin Biosynthesis in Tomato Fruits. Plant Communications 1, 100006.

Corso M, Perreau F, Mouille G, Lepiniec L. 2020. Specialized phenolic compounds in seeds: structures, functions, and regulations. Plant Science 296, 110471.

Dobin A, Davis CA, Schlesinger F, Drenkow J, Zaleski C, Jha S, Batut P, Chaisson M, Gingeras TR. 2013. STAR: Ultrafast universal RNA-seq aligner. Bioinformatics 29, 15–21.

Fallah AA, Sarmast E, Jafari T. 2020. Effect of dietary anthocyanins on biomarkers of glycemic control and glucose metabolism: A systematic review and meta-analysis of randomized clinical trials. Food Research International 137, 109379.

Fernandez-Pozo N, Menda N, Edwards JD, et al. 2015. The Sol Genomics Network (SGN)- from genotype to phenotype to breeding. Nucleic Acids Research 43, D1036–D1041.

Fu Z, Shang H, Jiang H, et al. 2020. Systematic Identification of the Light-quality Responding Anthocyanin Synthesis-related Transcripts in Petunia Petals. Horticultural Plant Journal 6, 428– 438.

Gao Y, Liu J, Chen Y, Tang H, Wang Y, He Y, Ou Y, Sun X, Wang S, Yao Y. 2018. Tomato SlAN11 regulates flavonoid biosynthesis and seed dormancy by interaction with bHLH proteins but not with MYB proteins. Horticulture Research 5, 1–18.

Gonzali S, Mazzucato A, Perata P. 2009. Purple as a tomato: towards high anthocyanin tomatoes. Trends in Plant Science 14, 237–241.

Gould KS, Dudle DA, Neufeld HS. 2010. Why some stems are red: Cauline anthocyanins shield photosystem II against high light stress. Journal of Experimental Botany 61, 2707–2717.

Grabherr MG, Haas BJ, Yassour M, et al. 2011. Full-length transcriptome assembly from RNA-Seq data without a reference genome. Nature Biotechnology 29, 644–652.

Gu Z, Eils R, Schlesner M. 2016. Complex heatmaps reveal patterns and correlations in multidimensional genomic data. Bioinformatics 32, 2847–2849.

Hardtke CS, Gohda K, Osterlund MT, Oyama T, Okada K, Deng XW. 2000. HY5 stability and activity in *Arabidopsis* is regulated by phosphorylation in its COP1 binding domain. EMBO Journal 19, 4997–5006.

Hichri I, Heppel SC, Pillet J, Léon C, Czemmel S, Delrot S, Lauvergeat V, Bogs J. 2010. The basic helix-loop-helix transcription factor MYC1 is involved in the regulation of the flavonoid biosynthesis pathway in grapevine. Molecular Plant 3, 509–523.

Hong Y, Tang X, Huang H, Zhang Y, Dai S. 2015. Transcriptomic analyses reveal species-specific light-induced anthocyanin biosynthesis in chrysanthemum. BMC Genomics 16, 1–18.

Houghton A, Appelhagen I, Martin C. 2021. Natural blues: Structure meets function in anthocyanins. Plants 10, 1–22.

Jiang M, Ren L, Lian H, Liu Y, Chen H. 2016. Novel insight into the mechanism underlying light-controlled anthocyanin accumulation in eggplant ( *Solanum melongena* L.). Plant Science 249, 46–58.

Jin J, Tian F, Yang D, Meng Y, Kong L, Luo J, Gao G. 2016. PlantTFDB 4. 0LJ: toward a central hub for transcription factors and regulatory interactions in plants. Nucleic Acids Research 45, 1040–1045.

Kanehisa M, Sato Y, Morishima K. 2016. BlastKOALA and GhostKOALA: KEGG Tools for Functional Characterization of Genome and Metagenome Sequences. Journal of Molecular Biology 428, 726–731.

Kim MJ, Kim P, Chen Y, Chen B, Yang J, Liu X, Kawabata S, Wang Y, Li Y. 2021. Blue and UV-B light synergistically induce anthocyanin accumulation by co-activating nitrate reductase gene expression in *Anthocyanin fruit* (*Aft*) tomato. Plant Biology 23, 210–220.

Larkin MA, Blackshields G, Brown NP, et al. 2007. Clustal W and Clustal X version 2.0. Bioinformatics 23, 2947–2948.

Levin I, Frankel P, Gilboa N, Tanny S, Lalazar A. 2003. The tomato dark green mutation is a novel allele of the tomato homolog of the *DEETIOLATED1* gene. Theor Appl Genet 106, 454– 460.

Li S, Dong Y, Li D, Shi S, Zhao N, Liao J, Liu Y, Chen H. 2024. Eggplant transcription factor SmMYB5 integrates jasmonate and light signaling during anthocyanin biosynthesis. Plant Physiology 194, 1139–1165.

Li YY, Mao K, Zhao C, Zhao XY, Zhang HL, Shu HR, Hao YJ. 2012. MdCOP1 ubiquitin E3 ligases interact with MdMYB1 to regulate light-induced anthocyanin biosynthesis and red fruit coloration in apple. Plant Physiology 160, 1011–1022.

Li J, Ren L, Gao Z, Jiang M, Liu Y, Zhou L, He Y, Chen H. 2017. Combined transcriptomic and proteomic analysis constructs a new model for light-induced anthocyanin biosynthesis in eggplant (*Solanum melongena* L.). Plant Cell and Environment 40, 3069–3087.

Liao Y, Smyth GK, Shi W. 2014. FeatureCounts: An efficient general purpose program for assigning sequence reads to genomic features. Bioinformatics 30, 923–930.

Liu CC, Chi C, Jin LJ, Zhu J, Yu JQ, Zhou YH. 2018a. The bZip transcription factor *HY5* mediates *CRY1a*-induced anthocyanin biosynthesis in tomato. Plant Cell and Environment 41, 1762–1775.

Liu Y, Tikunov Y, Schouten RE, Marcelis LFM, Visser RGF. 2018b. Anthocyanin Biosynthesis and Degradation Mechanisms in Solanaceous Vegetables: A Review. Frontiers in Chemistry 6, 52.

Liu C, Yao X, Li G, Huang L, Xie Z. 2020. Transcriptomic profiling of purple broccoli reveals light-induced anthocyanin biosynthetic signaling and structural genes. PeerJ 2020, 1–29.

Lloyd A, Brockman A, Aguirre L, Campbell A, Bean A, Cantero A, Gonzalez A. 2017. Advances in the MYB-bHLH-WD Repeat (MBW) pigment regulatory model: Addition of a WRKY factor and co-option of an anthocyanin MYB for betalain regulation. Plant and Cell Physiology 58, 1431–1441.

Love MI, Huber W, Anders S. 2014. Moderated estimation of fold change and dispersion for RNA-seq data with DESeq2. Genome Biology 15, 1–21.

Martin C, Butelli E, Petroni K, Tonelli C. 2011. How can research on plants contribute to promoting human health? Plant Cell 23, 1685–1699.

Meissner R, Jacobson Y, Melamed S, Levyatuv S, Shalev G, Ashri A, Elkind Y, Levy A. 1997. A new model system for tomato genetics., 1465–1472.

Mes PJ, Boches P, Myers JR, Durst R. 2008. Characterization of tomatoes expressing anthocyanin in the fruit. Journal of the American Society for Horticultural Science 133, 262–269.

Muraki I, Imamura F, Manson JE, Hu FB, Willett WC, Van Dam RM, Sun Q. 2013. Fruit consumption and risk of type 2 diabetes: Results from three prospective longitudinal cohort studies. BMJ (Online) 347, 1–15.

Panchal SK, John OD, Mathai ML, Brown L. 2022. Anthocyanins in Chronic Diseases: The Power of Purple. Nutrients 14, 1–30.

Pfaffl MW. 2001. A new mathematical model for relative quantification in real-time RT – PCR. Nucleic Acids Research 29, 16–21.

Podolec R, Ulm R. 2018. Photoreceptor-mediated regulation of the COP1/SPA E3 ubiquitin ligase. Current Opinion in Plant Biology 45, 18–25.

Povero G, Gonzali S, Bassolino L, Mazzucato A, Perata P. 2011. Transcriptional analysis in high-anthocyanin tomatoes reveals synergistic effect of *Aft* and *atv* genes. Journal of Plant Physiology 168, 270–279.

Qiu Z, Wang X, Gao J, Guo Y, Huang Z, Du Y. 2016. The tomato *Hoffman’s Anthocyaninless* gene encodes a bHLH transcription factor involved in anthocyanin biosynthesis that is developmentally regulated and induced by low temperatures. PLoS ONE 11, 1–22.

Qiu Z, Wang H, Li D, Yu B, Hui Q, Yan S, Huang Z, Cui X, Cao B. 2019. Identification of Candidate HY5-Dependent and -Independent Regulators of Anthocyanin Biosynthesis in Tomato. Plant and Cell Physiology 60, 643–656.

Quattrocchio F, Verweij W, Kroon A, Spelt C, Mol J, Koes R. 2006. PH4 of Petunia is an R2R3 MYB protein that activates vacuolar acidification through interactions with basic-helix-loop-helix transcription factors of the anthocyanin pathway. Plant Cell 18, 1274–1291.

Saijo Y, Sullivan JA, Wang H, Yang J, Shen Y, Rubio V, Ma L, Hoecker U, Deng XW. 2003. The COP1-SPA1 interaction defines a critical step in phytochrome A-mediated regulation of HY5 activity. Genes and Development 17, 2642–2647.

Sapir M, Oren-Shamir M, Ovadia R, et al. 2008. Molecular aspects of *Anthocyanin fruit* tomato in relation to *high pigment-1*. Journal of Heredity 99, 292–303.

Sestari I, Zsögön A, Rehder GG, Teixeira L de L, Hassimotto NMA, Purgatto E, Benedito VA, Peres LEP. 2014. Near-isogenic lines enhancing ascorbic acid, anthocyanin and carotenoid content in tomato (*Solanum lycopersicum* L. cv Micro-Tom) as a tool to produce nutrient-rich fruits. Scientia Horticulturae 175, 111–120.

Sivankalyani V, Feygenberg O, Diskin S, Wright B, Alkan N. 2016. Increased anthocyanin and flavonoids in mango fruit peel are associated with cold and pathogen resistance. Postharvest Biology and Technology 111, 132–139.

Sun C, Deng L, Du M, Zhao J, Chen Q, Huang T, Jiang H, Li CB, Li C. 2020. A Transcriptional Network Promotes Anthocyanin Biosynthesis in Tomato Flesh. Molecular Plant 13, 42–58.

Team RC. 2013. R: A language and environment for statistical computing. R Foundation for Statistical Computing, Vienna, Austria. http://www.R-project.org/, 201.

Wickham H. 2016. Data analysis. ggplot2. Springer, 189–201.

Wimalanathan K, Lawrence-Dill CJ. 2021. Gene Ontology Meta Annotator for Plants (GOMAP). Plant Methods 17, 1–14.

